# USA300 MRSA lineages persist on multiple body sites following infection

**DOI:** 10.1101/192096

**Authors:** Timothy D. Read, Robert A. Petit, Zachary Yin, Tuyaa Montgomery, Moira C. McNulty, Michael Z. David

## Abstract

**Abstract**

**BACKGROUND:** USA300 methicillin-resistant *Staphylococcus aureus* (MRSA) is a community- and hospital-acquired pathogen that frequently causes infections but also can survive on the human body asymptomatically as a part of the normal flora. We devised a comparative genomic strategy to track colonizing USA300 at different body sites after *S. aureus* infection.

**METHODS:** We sampled ST8 *S. aureus* from subjects at the site of a first known MRSA infection. Within 60 days of this infection and again 12 months later, each subject was tested for asymptomatic colonization in the nose, throat and perirectal region. 93 *S. aureus* strains underwent whole genome shotgun sequencing.

**RESULTS:** Genome sequencing revealed that 23 patients carried USA300 intra-subject lineages (ISLs), defined as having an index infection isolate (III) and closely related strains. Pairwise distance between strains in different ISLs was 48 to 162 single nucleotide polymorphisms (SNPs), whereas within the same ISL it was 0 to 26 SNPs. At the initial sampling time among 23 subjects, we isolated *S. aureus* from the nose, throat and perirectal sites from 15, 11 and 15 of them, respectively. Twelve months later we isolated *S. aureus* within the same ISL from 9 subjects, with 6, 3 and 3 strains from the nose, throat and perirectal area, respectively. The median time from initial acquisition of the *S. aureus* USA300 strains to culture of the index infection was estimated at 18 weeks. Strains in ISLs from the same subject differed in plasmid and prophage content, and contained deletions that removed the *mecA-*containing SCC*mec* and ACME regions. Five strains contained frameshift mutations in *agr* toxin-regulating genes. Persistence of an ISL was not associated with clinical or demographic subject characteristics.

**CONCLUSION:** Clonal lineages of USA300 may continue to colonize people at one or more anatomic sites up to a year after an initial infection and experience loss of the SCC*mec*, loss and gain of other mobile genetic elements, and mutations in the agr operon.

## Background

*Staphylococcus aureus* is one of the most common human bacterial pathogens, causing skin and soft tissue infections (SSTI), bacteremia, osteomyelitis and other disseminated infections [1]. While it can be deadly, *S. aureus* is more commonly a commensal species, living on the skin, the mucous membranes, and in the gut [2] of 25-50% of the human population [3]. Since the 1940s, the species has evolved resistance to each class of antibiotics introduced to treat infections [4]. Resistance to beta-lactam antibiotics due to the expression of penicillin-binding protein PBP2a, has been a particularly vexing problem. Strains that express PBP2a, known as methicillin-resistant *S. aureus* (MRSA) have proliferated since being first reported in 1961 in the United Kingdom [5,6]. PBP2a is coded by the *mecA* gene, which is carried in a mobile genetic element called SCCC*mec*. Initially reported in just one MLST (multilocus sequence type) background, ST250, in the last 55 years, SCC*mec* cassettes have spread to many sequence types [7].

Previous studies have demonstrated distinct differences in fitness and virulence among genetic backgrounds of MRSA [8–10]. One genotype of MRSA, USA300, emerged in the United States (US) in the 1990s and spread rapidly in the community to become the leading cause of SSTIs in US urban emergency departments by 2004 [11]. USA300, which has also spread to many other parts of the world [12,13], is MLST ST8 and usually carries SCC*mec* type IV, the phage-encoded Panton-Valentine leukocidin (PVL), and the arginine catabolic mobile element (ACME) [14]. The genetic factors underlying the meteoric success of this strain type are not completely understood but it is likely that enhanced virulence promoting transmission, tolerance of the human body to its presence as a commensal, and an apparently limited fitness cost of its antibiotic resistance [15] are important. Only a few studies have suggested that USA300 can stably colonize patients for more than 6 months, but these have been limited by using genotyping methods that lack the genetic resolution to distinguish between persistent colonization by a strain and acquisition of a new colonizing strain [16]. However, two recent studies have used whole genome sequencing to demonstrate the persistence of single USA300 clones to cause recurrent infections in individuals [17,18].

There have now been a number of comparative genomics studies on USA300 MRSA. These have included studies using the fine scale resolving power of genomics to characterize the global population structure [13,19–21] and transmission patterns between patients [22,23], within households [24–26] and in recurrent SSTIs [17]. In this work we explored the diversity of USA300 on the bodies of subjects during a year of surveillance after an initial, clinically significant USA300 MRSA infection. In individual subjects within a prospective cohort, we set out to determine whether a single USA300 clonal lineage persisted after an infection, whether it persisted on or spread among three body sites, and how the genome changed in the clonal lineage during a year of observation.

## Methods

### Strain Isolation and Phenotypic Testing

Patients at the University of Chicago Medical Center (UCMC) with a first known MRSA infection at any anatomic site were enrolled in a longitudinal study of risk factors for recurrent MRSA infections between in April 2012 and February 2014. Subjects underwent an enrollment visit within 60 days of culture of their initial infection (time 1) when they were tested for *S. aureus* colonization of the nares, throat, and perirectal region. Subjects returned approximately one year later and were again cultured at the same three sites (time 2). A medical record review was performed to determine the site of the initial MRSA infection and to determine if a true infection was present [27], the therapy used to treat infection, demographics, and comorbidities. Each subject was administered a detailed survey to assess risk factors for MRSA infection. All *S. aureus* isolates, including the initial infection isolate (III) and up to a single isolate obtained from each colonization culture, were frozen at -80C. All of these isolates underwent genotyping by MLST, PCR for the presence of the PVL genes, and, for MRSA isolates, SCC*mec* typing.

For the present study, all subjects who had an ST8 MRSA III isolate and one or more ST8 *S. aureus* isolates obtained from colonization cultures at one or both colonization culture visits were included (n=31/169, 18.3%). Two of the III samples were lost, leaving 29 III from 31 patients. 93 isolates from these 31 subjects underwent genomic preps and Illumina sequencing. All strains produced hemolytic clear zones on blood agar plates.

**Illumina Sequencing**

Genomic preps were prepared with the DNeasy Blood and Tissue Kit (Qiagen, Germantown, MD). Libraries for sequencing 93 strains were constructed using Nextera technology (Illumina, Inc., San Diego, CA). Libraries were sequenced on an Illumina HiSeq instrument using either 100 nt or 125 nt paired-end protocols.

### Sequence Data Analysis

Only two strains had less than 100x predicted genome coverage (assuming an average 2.9 Mbp). For consistency, all other genomes were downsampled to 100x coverage before assembly. All but two strains (both III) were found to be ST8 by genome-based MLST. These two outlying strains were not included in subsequent analysis. FASTQ sequencing output files from strains with more than 100x coverage were downsampled using a custom script (https://gist.github.com/rpetit3/9c623454758c9885bf81d269e3453b76) based on the seqtk toolkit (https://github.com/lh3/seqtk). Strains were assembled de novo using SPAdes [28] and annotated using PROKKA [29]. Antibiotic resistance phenotypes were predicted for each strain based on the methods of Gordon et al. [30]. SnpEff 4.2 [31] was used to annotate SNPs in the VCF output of the parsnp alignment based on USA300_TCH1516 coordinates. SNPs were mapped back on the shortened alignment using R BioStrings [32]. USA300_TCH1516 proteins were mapped to their orthologs in the *S. aureus* type strain NCTC8325 [33] using BLASTP, and gene ontology enrichment was performed using PANTHER [34] in December 2016.

A whole genome alignment of all the newly sequenced USA300 genomes was performed using parsnp v1.2 [35] against the closed reference genome USA300_TCH1516 (NCBI refseq accession NC_010079.1). The SNIPPY pipeline [https://github.com/tseemann/snippy] was used to cross-validate SNPs predicted by parsnp and to identify frameshifts and small indels. Plasmid content was found to be variable between strains (discussed below), and therefore not useful for comparisons across the whole dataset. Two ST8 strains fell outside of the USA300 subgroup and were therefore excluded from the rest of the study, leaving a total of 89 strains. Due to these exclusions, 2 patients were not represented by any samples and we therefore concentrated our analysis on the 29 remaining patients.

The concatenated common aligned portion (2,471,416 of the 2,872,915 bp [86%] USA300_TCH1516 reference) was used as the input for the recombination detection program ClonalFrameML v1.0-19-g9488a80 [36], along with a maximum likelihood guide tree produced by RAxML [37] v7.2.8. We removed 10,884 bp from the alignment identified by ClonalFrameML [36] as possibly of recombinant origin and then used as input for a maximum likelihood phylogeny under the GTRGAMMA model with 100 bootstraps using RAxML. Noisy (v1.5.12) [38] was used to scan for potentially homoplasious sites in the alignment. The sequences of the inner nodes of the phylogeny were reconstructed using FASTML [39], based on the post- ClonalFrameML trimmed alignment. The R ape [40] dna.dist function was used to find the number of SNPs between leaves and ancestral nodes of the tree based on Hamming distance.

To identify integrated prophages, we aligned PCR primer sequences from seven diverse classes of *S. aureus Siphovirus* integrase [41] using blastn (version 2.3.0), with the ‘blastn-short’ task option. To identify plasmid contigs, we aligned against a database of 230 complete *S. aureus* plasmids downloaded from NCBI using the blastn ‘megablast’ option. The contigs were also aligned against sequences from a Gram positive plasmid replicon typing scheme [42], identifying plasmid replicon [42] matches with BLASTN (task = blastn-short, cutoff = 1 mismatch or gap, match of 19 bases or more). Visualization of the assemblies mapped by BLASTN against the reference chromosome was produced by the CGView Comparison Tool (CCT) [43].

Raw sequence data were deposited at NCBI SRA under bioproject accession PRJNA388087.

### Association of Subject Characteristics with Persistence of an Intra-Subject Lineage (ISL)

We performed univariate testing (chi square, Fisher exact, Wilcoxon rank-sum or t-tests) to determine if subject characteristics (see list in supplemental data) were associated with colonization with an ISL (see discussion of ISL below) after III at times 1 and 2. We used the Bonferroni correction to compensate for multiple comparisons.

## Results

### Colonizing ST8 *S. aureus* Strains Can Be Isolated 12 Months Following Initial Infection

We recruited an initial study cohort of 169 patients, among whom 29 presented with a ST8 MRSA infection and were subsequently found to have at least a single ST8 *S. aureus* isolate obtained from a colonization culture at one or more of the 3 tested body sites at time 1 (enrollment) or time 2 (12 months). Study subjects received 0-6 antibiotic drugs during treatment for the index infection (Additional File 1). *S. aureus* III from the site of infection were retained prospectively for further study and underwent whole genome sequencing. Nose, throat and perirectal body sites were cultured at time 1, and *S. aureus* was isolated from 16, 15 and 17 patients, respectively. Approximately 12 months after the initial visit (time 2), we isolated *S. aureus* from 6, 3 and 4 subjects from nose, throat and perirectal sites, respectively (Table 1 and Additional File 1). While all III isolates were obtained from suspected sites of infection, on review of the medical record, 2 were found to be obtained from sites of colonization and did not require treatment.

**Table 1.** Demographic and Clinical Information. An extended version of this table listing additional variables collected is in the supplemental data.

| Patient | Ill | ISL | strain 1* | strain 2* | Sex | Age | Description |
| --- | --- | --- | --- | --- | --- | --- | --- |

| ID |  |  |  |  |  | (years)** |  |
| --- | --- | --- | --- | --- | --- | --- | --- |
| 1 | y | y | y | y | female | 55 | respiratory |
| 2 | y | y | y | n | female | 1 | perirectal abscess |
| 3 | y | y | y | n | female | 0 | neck abscess |
| 4 | y | y | n | y | male | 0 | right buttock abscess |
| 5 | y | y | y | n | female | 13 | draining boil |
| 6 | y | y | y | n | female | 56 | abdominal wall abscess |
| 7 | y | y | y | n | male | 10 | left groin abscess |
| 8 | y | y | n | y | male | 0 | pleural fluid culture |
| 9 | y | y | y | y | male | 48 | right ear drainage |
| 10 | y | n | n | n | female | 35 | abscess right shoulder |
| 11 | y | y | y | y | male | 9 | left ear drainage |
| 12 | y | y | y | n | male | 3 | abscess near anus |
| 13 | y | y | y | y | female | 1 | left labial cellulitis and abscess |
| 14 | y | y | y | n | female | 1 | left labial abscess |
| 15 | y | y | y | n | female | 44 | drainage from psoriatic plaque |
| 16 | y | y | y | n | female | 52 | foot wound |
| 17 | y | n | n | n | male | 1 | left inguinal abscess |
| 18 | y | y | y | y | female | 59 | sputum culture |
| 19 | y | y | y | y | female | 66 | left leg wound<br>drainage |
| 20 | y | y | n | y | female | 0 | wound drainage neck,<br>axilla, groin |
| 21 | y | n | n | n | male | 1 | buttock abscess |
| 22 | y | y | y | n | male | 0 | culture dorsum right<br>hand |
| 23 | y | n | n | n | female | 43 | right midface abscess |
| 24 | y | y | y | n | female | 31 | left upper arm<br>abscess |
| 25 | y | y | y | n | male | 6 | wound drainage,<br>scrotal abscess |
| 26 | y | n | n | n | male | 36 | unknown |
| 27 | y | y | y | n | female | 36 | wound drainage leg<br>abscess |
| 28 | y | y | y | n | male | 29 | wound drainage chin<br>abscess |
| 29 | n | n | n | n | male | 0 | respiratory |

\*whether or not a USA300 strain within an ISL was isolated at visit 1 or 2 (one year later) \*\*at
initial visit

### Phylogeny of USA300 Isolates from Chicago Patients Reflect History of the Local Epidemic

From the alignment of the 89 strains from 29 patients against reference USA300_TCH1516 isolated from a pediatric patient at the Texas Children’s Hospital before 2007 [44], we extracted a core region of the USA300 chromosome of 2,471 kbp (86%). The 14% of the chromosome excluded because it was missing in at least one of the genomes included SCC*mec*, ACME, the SaP15 pathogenicity islands, and the SLT and βC prophages [44]. Other regions not part of the core alignment were plasmids, rRNA operons, transposons, portions of membrane proteins and non-genic repeats.

The strain phylogeny (Figure 1) was composed of clades of leaf nodes with short branches near the tips, mostly representing isolates from individual patients, separated by relatively long branches in the middle section. In the deepest parts of the tree, branches were shorter and had lower bootstrap confidence. This pattern, seen in other genome-based studies of USA300 [24,25] reflected the history of the epidemic [45]. USA300 spread rapidly after its introduction in the 1990s to US urban areas such as Chicago and achieved a large population size. USA300 then persisted in the community, causing sporadic clinically significant infections. By rooting the tree with the TCH1516 reference, a group of strains with mutations in *gyrA* and *grlA* predicted to confer ciprofloxacin resistance fell into a late-branching subgroup. The later development of resistance to fluoroquinolones was in line with results from analysis of the global population of USA300 [13,20,24,25].

**Figure 1.**
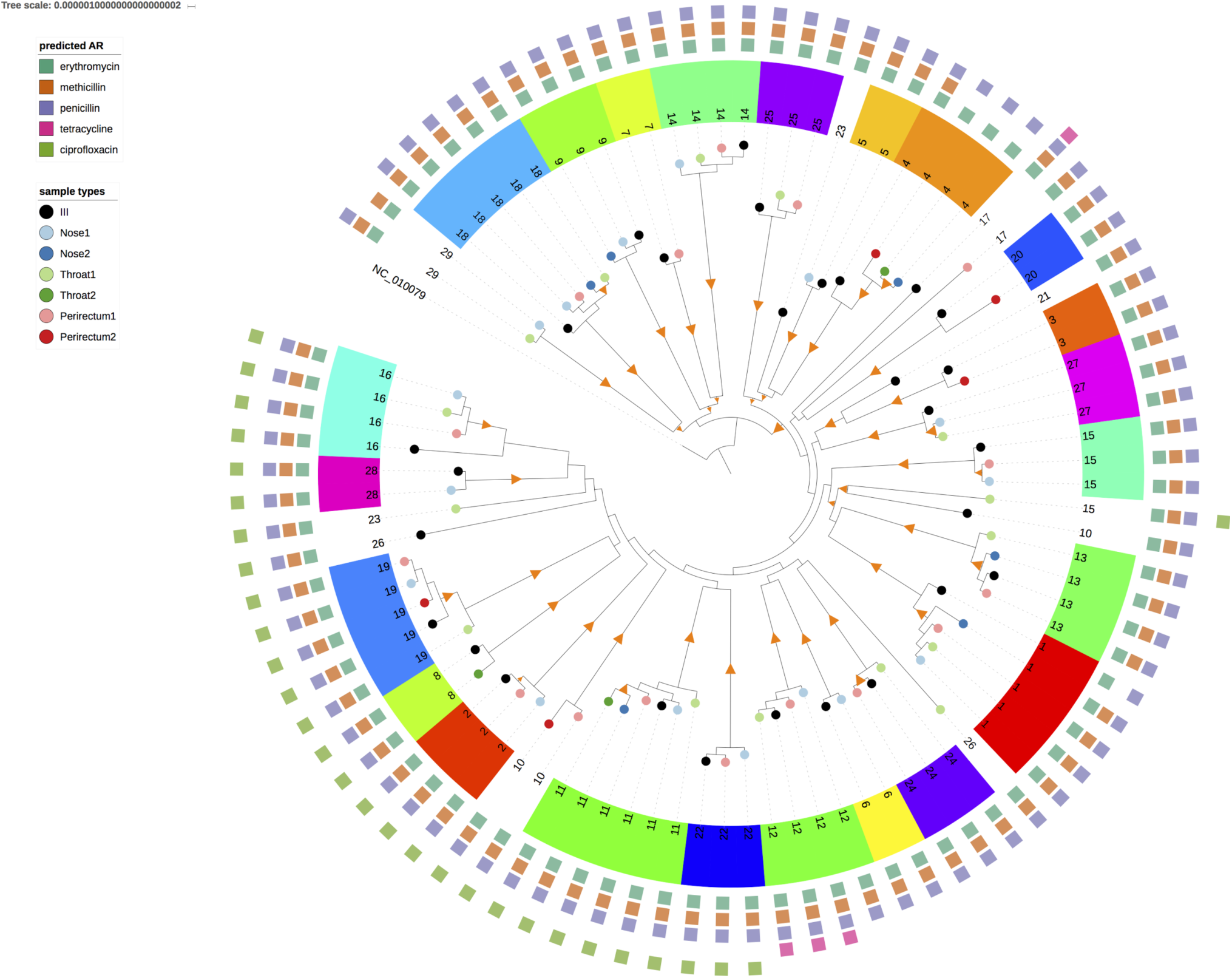
Maximum Likelihood tree of USA300 strains sequenced in the project. Orange triangles are branches with > 80-100% bootstrap support. Tips of trees are labelled by site of isolation (black = index infection isolate (III), green = throat, blue = nose, red = perirectal). Light colors are from sampling time 1, dark colors from sampling time 2. A color code and patient number was assigned to each patient. The squares on the outer ring represent antimicrobial class resistance predicted by the genome sequence. The figure was drawn using iTOL software [70].

### USA300 Strains Form “Intra-Subject Lineages” (ISLs) Present on Multiple Anatomic Sites of a Subject’s Body

As mentioned above, in most cases, the strains isolated from one subject clustered into a monophyletic group with relatively short branch lengths. We termed these groups ISLs, and we inferred that all strains within them derived from a single recent introduction to the subject’s body. Based on the phylogeny we removed six strains from ISLs because they were obviously phylogenetically unrelated to the other strains in the ISL. Five subjects (10,15,17,23,26) had colonizing strains from a different ISL than the III strain, suggesting infection with more than one USA300 lineage. After removing these non-conforming strains, we were left with 23 subjects with an ISL that included an III isolate and at least one other body site isolate. For 14 of these ISLs, the III branched first (with > 80% bootstrap support) from the predicted Most Recent Common Ancestor (MRCA) relative to the other nodes, as would be expected in a scenario where the infection occurred soon after an initial introduction of a strain that subsequently spread to other body sites. In three of the ISLs in which the III branched first, there was at least one other strain, obtained from a colonization culture, with identical sequence (no SNPs). In the other nine ISLs a strain other than the III branched first, suggestive of a MRSA infection following earlier colonization.

Of the 23 patients with a USA300 ISL, 15, 11, and 15 (50%, 39% and 54%) at the time 1 (enrollment) visit had an an isolate from the same ISL in the perirectal area, throat and nose, respectively. Following up one year later (i.e., time 2), 6 (21%) patients still had a strain from the same ISL in the nose, and 3 (11%) each in the throat in the perirectal area. Strains were isolated at time 2 in 9 of the of original 23 ISLs (Table 1). Fourteen subjects had more than one body site colonized by a strain from within an ISL at enrollment (time 1); only 2 subjects were colonized at more than one site with an ISL a year later.

### Lack of Association Between Subject Characteristics and Persistence of an ISL

In univariate analyses, we found that no tested subject characteristics, including age, sex, type of insurance coverage, race, treatment of the index infection with clindamycin, with vancomycin or with trimethoprim-sulfamethoxazole, or pet exposure (see complete list in supplemental data) were significantly associated with colonization with an ISL after III at times 1 and 2 after Bonferroni correction.

### Genetic Distance Between and Within ISLs

For each strain we calculated the number of SNPs between every other strain in another ISL and compared it to the number of SNPs within an ISL (Figure 2). Outside of the one possible example of direct transmission between patient 3 and 30 (Figure 1), strains in different patients had from 48 to 162 SNP differences, with a median of 106. The range for strains in the same ISL was 0 to 26 with a median of 5.

**Figure 2.**
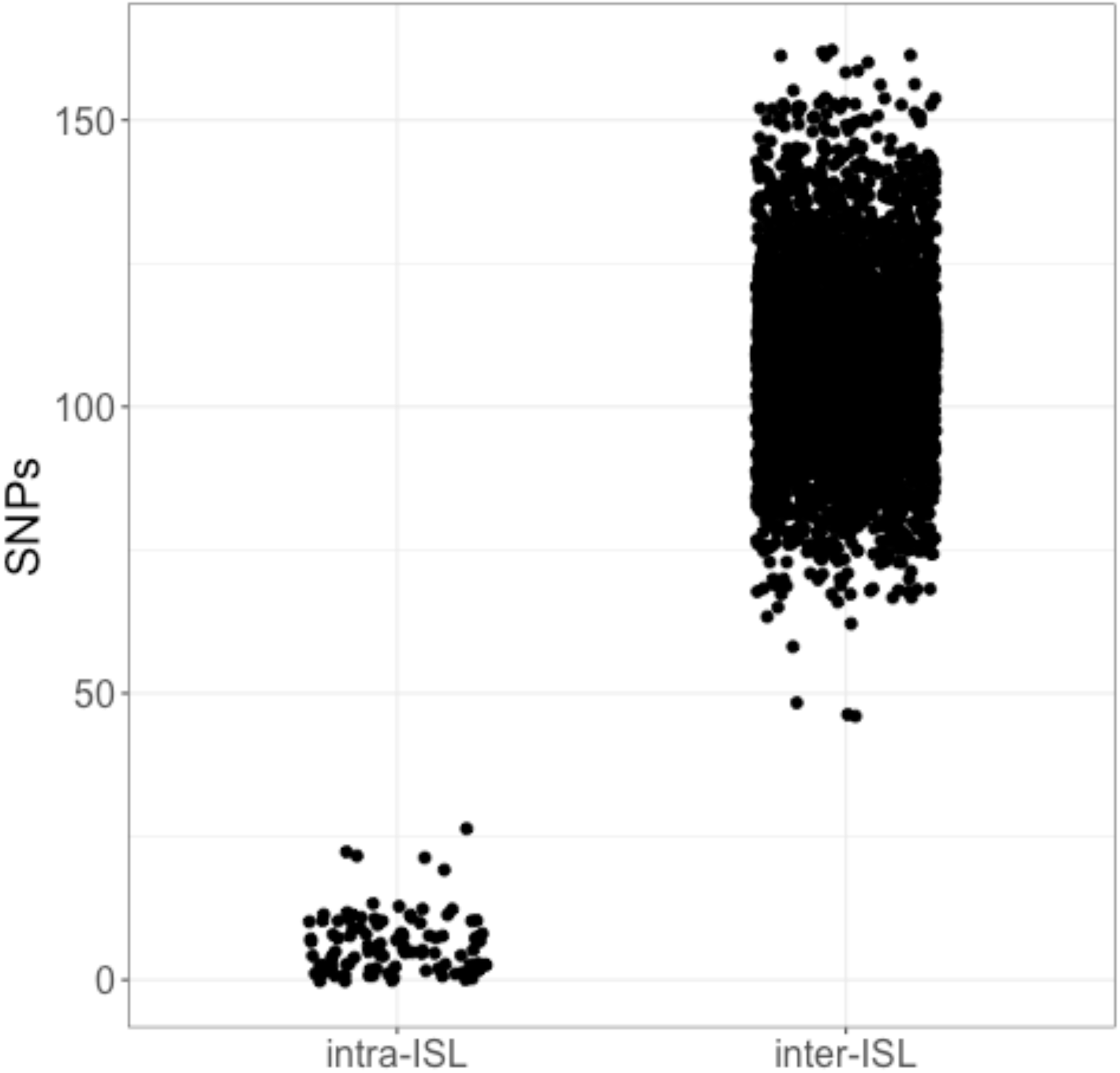
Distribution of the number of SNPs separating strains within the same ISL (intra- subject lineage) compared to other strains.

The intra-ISL relationships were further analyzed by measuring the number of SNPs between each strain and the III (Figure 3). As expected, within host analysis showed that there were a significantly greater number of SNPs between the III and strains from time 2 (range 2-26: median 10) compared with time 1 (range 0-12: median 3) (t-test, P < 7.8e-7). However, there was no significant pattern to the number of SNPs differing in the III and each of the three tested body sites (ANOVA p = 0.8). The ranges for each body site were: nose, 0-22 (median 4.5), throat, 0-19 (median 7) and perirectal 0-16 (median 3.5).

**Figure 3.**
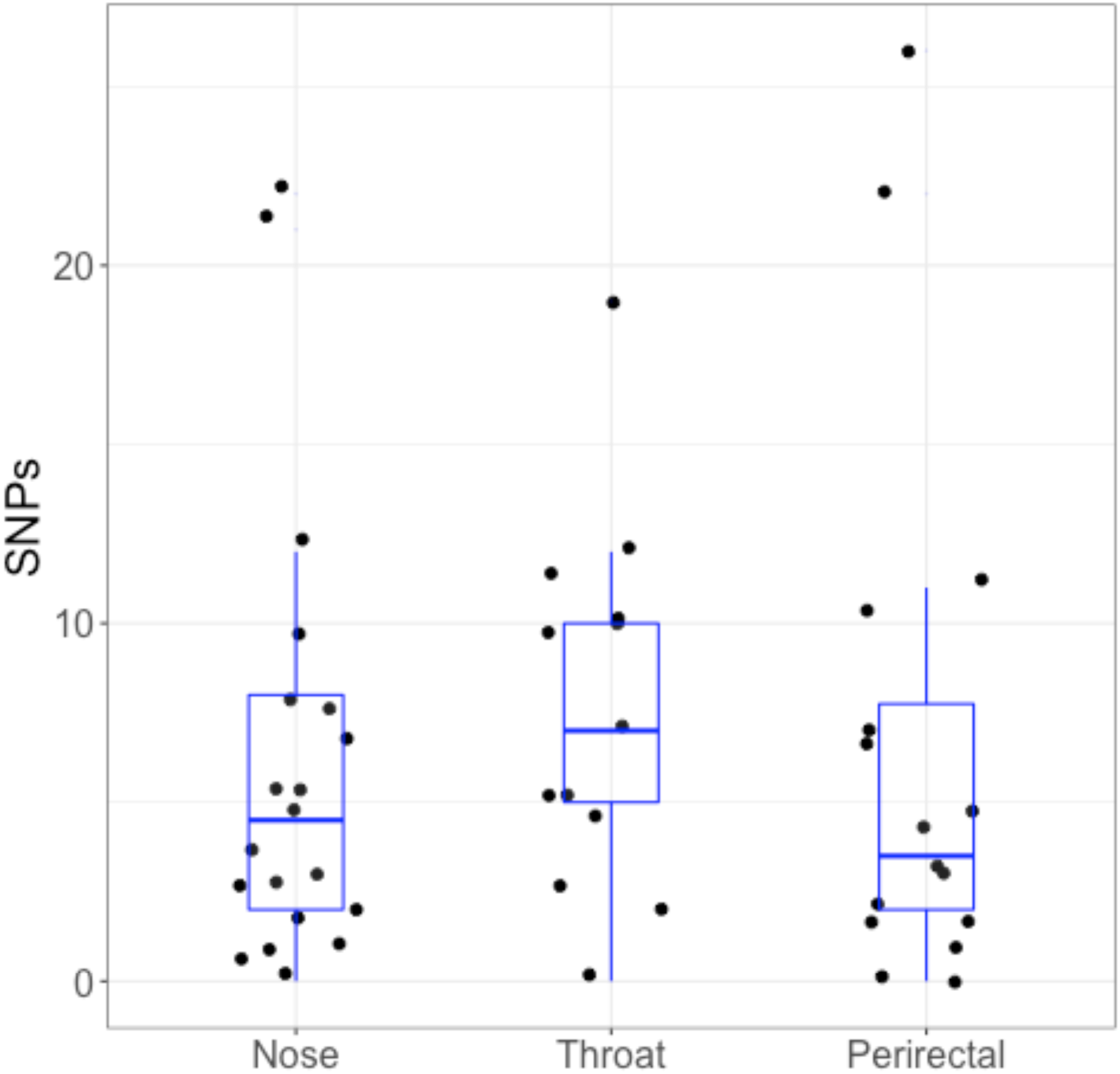
Number of SNPs separating the III and strains within the same ISL at different sites. Horizontal line in the box show the median, and box hinges are the 25th and 75th quartile. The upper whisker extends from the hinge to the largest value no further than 1.5 * inter-quartile range (IQR) from the hinge. The lower whisker extends from the hinge to the smallest value at most 1.5 * IQR of the hinge.

We used FASTML [39] to reconstruct the sequence of internal nodes of the phylogeny and then found the number of SNPs differing between each strain of an ISL and its MRCA. There was a significantly greater number of SNPs from the MRCA at time 1 (median = 2) compared with time 2 (median = 9.5) (t-test, p < 7.7 e-13) (Figure 4A).

**Figure 4.**
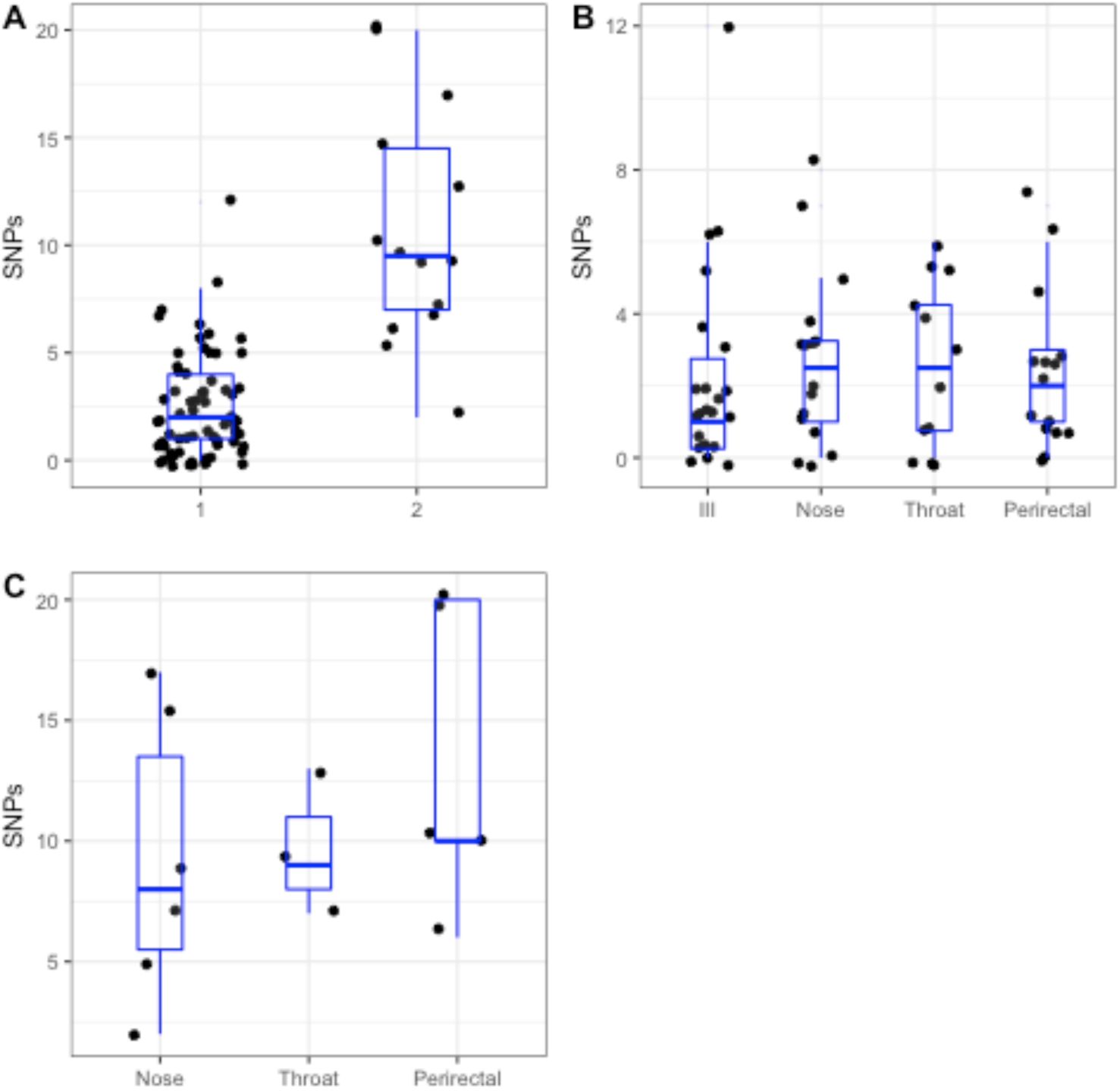
Number of SNPs between the MRCA and strains at each sampling time point: 4A, aggregated for each time point. Sampling time point 2 was approximately 1 year after time point 1. 4B, at time point 1 between the MRCA and each body site. 4C, at time point 2 between the MRCA and each body site.

Multiple genome-based studies have estimated the *S. aureus* mutation rate to be between 1.3- 3.3 x 10-^6^ SNPs per genome per year [24,46–49]: approximately 1 SNP every 9 weeks [2]. Based on this estimate, the median time from initial acquisition of the *S. aureus* USA300 strains to the clinical culture would have been about 18 weeks. There was no significant difference in the number of SNPs between the III or other body sites at time 1 (ANOVA, p = 0.95), or at the three body sites sampled at time 2 (ANOVA, p = 0.49) (Figure 4B and C).

### Shifts in Mobile Genetic Element Composition Within ISLs

In *S. aureus* changes in accessory gene content mediated by prophage and genome island insertional and plasmid transfer are important mechanisms for genome evolution. However, tracking these changes is problematic using short read shotgun data. Loss or gain of a single element might affect dozens of genes at once, and plasmids and prophages undergo frequent recombination and share common sequences, making identification difficult [50]. Nevertheless we were able to use previously developed typing schemes to gain insights into changes in mobile element composition during colonization of the studied subjects.

For prophages, we used the scheme of Goerke et al., who defined 7 distinct sequence types of *S. aureus* integrase (Sa1-7) [41]. Presence of the each class was called based on BLAST matches of at least one of the integrase-specific primers. Most strains had Sa2 (86/89; 96%) and Sa3 (87/89; 98%) family prophages. These correspond to the ϕSa2 (ϕSLT) and ϕSa3 (ϕBC) prophages in the USA300_TCH1516 reference genome, respectively [44]. In addition, 40/89 (45%) strains had a Sa5 integrase gene. Seven strains (8%), all from patients 13 and 15, carried an Sa1 integrase. The patterns of prophage carriage mirror those described in USA300 by Jamrozy et al. [20], with the exception that we did not identify the rarer ϕSa7 prophages in our sample. We noted four occurrences of prophage composition change within an ISL (Additional File 2). These were the apparent loss of ϕSa3 in the III of subject 18 and the first throat culture of subject 15 and the apparent gain of ϕSa5 in the first perirectal isolate of subject 10 and the III of subject 16 (although the III may represent a separate infection event from other strains isolated from subject 16 based on its deep branching).

To examine plasmid content we used the combination of BLAST alignments against DNA sequences from the Gram positive replicon typing scheme of Lozano et al. [42] and 344 complete *Staphylococcus spp*. plasmid sequences downloaded from NCBI (Additional File 2). All genomes contained the common backbone region of the 27 kb antibiotic-resistance plasmid of TCH1516, pUSA300HOUMR (NC_010063) [44]. 67/89 (72%) strains matched against > 25 kb of the plasmid, the rest subsequences of the plasmid as small as 13kb that all contained the pN315-like replicon [51]. Eight strains contained most of the 37 kb pUSA03 [52] multidrug resistant conjugative plasmid sequence; notably, the *ileS* mupirocin resistance gene was deleted. In the Jamrozy et al. study, in contrast to our findings, 146/154 (95%) USA300 strains from SSTIs carried the pUSA300HOUMR backbone plasmids and 5% (8/154) carried pUSA03- like plasmids. Families of smaller (< 10 kb) plasmids were present in some of our strains. The cryptic pUSA01 plasmid was found in 59 (66%) strains; pC194-like plasmids [53] were found in 6; 4 strains carried a plasmid with tetracycline resistance gene t*etK* similar to the pUSA02 plasmid [52] (NC_007791); and there were 3 and 1 strains with plasmid replicons of the pUB110 [54] and pWG745 [55] families, respectively.

Eight ISLs had strains with different plasmid content (patients 1,3,7,11,12,16,17; Figure 1). Plasmid was not linked to a particular body site or sampling time point. In subject 11,16 strains appeared to have acquired a pC194-like plasmid during colonization. In subject 12, the perirectal isolate at the time 1 (enrollment) was apparently cured of two plasmids (pUSA02, pUSA03) carried by all the other strains.

### Predicted Antibiotic Resistance Variation within ISLs

We noted that strains within several ISLs varied in their antibiotic resistance profile, based on the genomic predictions using the method of Gordon et al. [30]. The first throat isolate of subject 15 acquired ciprofloxacin resistance through *gyrA/grlA* mutations. This isolate was not part of the same ISL as the III, nose and perirectal isolates from subject 15 (Figure 1), and therefore it was not possible to determine whether the mutations occurred during colonization of this subject or earlier. Subject 15 was an outpatient who received only clindamycin during treatment of the index infection.

Two subjects had strains containing a 4.5 kb plasmid > 99% similar to pT181 [56] (NC_001371.1) and pUSA02[52] (NC_007791), with the *tet(K)* tetracycline resistance determinant. One of these was the III of subject 17, which was in a different ISL from the other strains isolated from this person (Figure 1). From subject 12, the III and 2 other strains contained the *tetK* plasmid, but the nose strain did not. The most parsimonious explanation for the phylogenetic pattern was that the strain originally infecting the subject was tetracycline resistant and the nose isolate had been cured of the plasmid. Neither subject 12 nor 17 were recorded as being treated with a tetracycline.

In subject 30, who was not treated with a β-lactam antibiotic for the index infection, the nose isolate had lost its *blaZ* gene owing to a deletion. The throat isolate from subject 30 was was identical in its core chromosome sequence and had a plasmid with no deletion. In two subjects, strains switched from MRSA to MSSA because of deletion of the SCC*mec* cassette, which included the *mecA* gene. On subject 4 the III was MRSA but all three strains isolated at time 2 (the 1-year visit) were MSSA. For subject 1, the situation was reversed. The III was MSSA, but the other strains isolated were all MRSA. The most parsimonious explanation was loss of the genes in the ancestor of the III shortly after colonization of the subject. Neither subject 1 nor 4 was treated with a β-lactam antibiotic for the index infection.

### SNP Patterns Do Not Show Signatures of Strong Positive Selection within ISLs

Of the 1,882 SNPs in core regions, 1,494 (79%) were in genes and of these, 1,101 (73%) caused non-synonymous or premature termination changes. This high percentage of mutations affecting amino acid sequences is typical when comparing closely related bacteria before purifying selection has time to operate [57] and has been reported previously in studies comparing USA300 strains [17,20]. The non-synonymous changes fell over 787 genes. Using a Poisson test, we did not find a significant deviation from the number of mutations that would be expected based on the gene length, if the mutations were occurring randomly across the genome. The gene with most changes was USA300HOU_1372, encoding an extracellular binding protein precursor (*ebp*) with 27 mutations; this protein is also the largest in the genome (10,421 amino acids). USA300HOU_0192 encoding a non-ribosomal peptide synthase (2397 aa), had the second most changes with 7 mutations. Most mutations were only found in one strain and more than 95% were found in 7 or fewer genomes. This was a result of the sampling strategy employed in the study, where a maximum of seven strains was taken from the same subject. Mutations that occurred frequently were those found only in the reference (and hence common to all 89 strains sequenced here), in deeper branches of the tree, and those causing ciprofloxacin resistance.

We looked to see if there were any functional differences between SNPs that occurred only in a certain body site, as it is possible that there was tissue dependent selection that may act over multiple genes linked by common phenotype. We found 50, 190, 187 and 246 SNPs unique to nose, perirectal area, throat and III, respectively (the lower number for nose was because there were no isolates from this site on isolated branches of the phylogeny (Figure 1). Using PANTHER [34] we looked for pathways in the Gene Ontology enriched for genes affected by terminations and non-synonymous mutations at each site but found no significant results after multiple hypothesis correction.

### Deletions in Agr and ACME/SCC*mec*/ϕSa2 Deletions Within ISLs

We found that chromosomal indels reshaped virulence and resistance elements in the III and colonization strains. ACME is a 31 kb mobile genetic element that has been suggested to be important for the success of North American USA300 strains [58] (although it absent from South American strains) [19]. Eleven strains in this study were found to have a deleted ACME and one partially deleted. In 6 subjects, the ACME element appears to have been lost by strains within the ISL (assuming the MRCA of the ISL was ACME+). In subject 25, who was not treated with any antibiotic for the index infection, all strains were deleted in their ACME element and also lost their flanking SCC*mec* cassette, making them MSSA. Two strains isolated from the perirectal region of subject 20 were missing the ϕSa2 prophage carrying the PVL *lukSF* genes. These accessory genes are postulated to be important for SSTIs caused by USA300 (discussed by Otto, 2013 [59]).

We also noted that 5 strains contained frameshift mutations in the key *agr* regulatory locus that would theoretically render them less capable of causing toxin-mediated tissue damage [60,61] (Additional File 2). However, these strains and all others isolated for this study were hemolytic when grown on blood agar plates, suggesting that these mutations did not completely disrupt the system. Two strains from subject 26 (in the III and first throat isolate) that were not part of the same ISL, had frameshift mutations in different portions of the *agrC* gene. Three of six strains from subject 11 isolated at both timepoints in all body sites carried the same frameshift in the *agrA* gene encoding the regulatory protein, illustrating that the *agr* was not necessary for long-term survival on patients and also showing that *S. aureus* strains with this mutation had migrated between body sites.

## Discussion

Assessing adult subjects with persistence of USA300 MRSA asymptomatic colonization after a clinically confirmed or suspected USA300 infection, we demonstrated that a single ISL was responsible for this persistence. Specifically, we showed that *S. aureus* ISLs, defined as having < 30 SNPs over 86% of the chromosome from the CA after about a year, colonized a patient over an extended time period. This cutoff falls in line with other recent studies. Young et al. [62] sequenced 68 nasal and blood isolates from a MSSA clone in one patient over six months, finding a total of 30 SNPs in all isolates. Sabat et al. reported that there were a maximum of 8 SNPs difference between USA300 isolated at the same time from different body sites of patients during a USA300 outbreak in Surinam [23]. Our finding here of stable colonization with a single USA300 lineage for a year or more is also in line with the result of the shotgun metagenomic study of Oh et al. [63], who found that strain-specific SNPs for another another human colonizing *Staphylococcus* species, *S. epidermidis*, could be recovered at the same skin sites at sampling times 1-2 years apart, suggesting temporal stability.

In most cases, strains isolated from the three tested body sites belonged to the same ISL as the isolate obtained from the index infection (i.e., the III), collected prior to the first test for colonization. We also demonstrated that the stable ISL clonal lineage had arrived on the patient recently before their enrollment in the study, on average within the previous few months (median of 2 SNPs from the MRCA to the first isolates; Figure 4A). This short time period from introduction to the index infection may be a feature of the high virulence of USA300 [59], and it might be interesting to compare to other *S. aureus* strain types. When the same patient was sampled a year later (time 2), the chance that an ST8 *S. aureus* strain could still be isolated was lower, but the ST8 isolates that were obtained at time point 2 were almost all from the same ISL.

Phylogenetic analysis suggested that in 13/23 ISL the III was either first branching or at least identical to another first branching strain. This would support the hypothesis that later colonizing strains derived from the ancestor of the III. In some cases, the tree supported an alternative scenario, in which USA300 spread to different body sites *before* the index infection was reported. Some caution must be used in interpreting trees from a small number of mutations, however, and these patterns will need to be confirmed by repeating this sampling approach with a larger number of patients. Nevertheless, our findings do raise important questions about the anatomic distribution of MRSA before and after a clinically significant infection. MRSA may have a widespread anatomic distribution prior to onset of an infection. By comparing the positions on the ISL phylogeny of strains from the first and second time point for colonization testing, it is possible that strains may have moved between or among body sites rather than just remaining diversifying in place. These results also suggested that the nose, throat and perirectal area were approximately equally efficient at maintaining *S. aureus* colonization, potentially challenging the enduring belief that the primary human site of long-term *S. aureus* colonization is the anterior nares [64]. Further studies are necessary to confirm this suggestion.

We did not identify a specific gene or locus that acquired an unusual number of SNPs suggestive of positive selection. With the number of genomes in our study, we had limited power to detect parallel mutations, especially those that may have been associated with adaptation specifically to the anatomic niches of the nose, throat or perirectal region. We did, however, encounter a significant number of genetic changes associated with movements of mobile elements during the history of colonization, including phage and plasmid gain and loss and loss of ACME and SCC*mec* elements. The latter two are particularly significant as they are considered central to the success of the USA300 MRSA clone, although naturally occurring deleted strains are not infrequently isolated [13,20,65]. We also encountered *agr* mutant strains, as have others who have studied *S. aureus* colonization [17,60,66]. These mutants may be evolutionary dead ends if transmission to new hosts preferentially selects for ACME+ SCC*mec*+ and agr+ strains.

This study had certain limitations. With limited resources we were constrained to sampling one colony at each site/time point but as sequencing costs fall it will be important to take multiple colonies to understand the extent of within strain variation on a single host at a single time point [67]. Recent studies have shown that rare variants within population can be important in tracking transmission among hosts [68,69]. It would also be invaluable to compare the *S. aureus* strain to the background skin microbiome in order to start to approach questions about the source of selective pressure to lose or gain genetic elements such as SCC*mec* and selective pressure leading to persistence or loss of *S. aureus* colonization.

Additional studies may help answer important questions related to the evolution of *S. aureus* during colonization, transmission and disease that could positively impact health outcomes. We don’t know how long an ISL persists on individuals and how effective various classes of systemic antibiotics are in reducing colonization. We know that individual clones persist for years in households but there may be expansion of a reservoir with frequent transfer of strains between individuals and fomites [24,25]. Is the site of long term colonization always the nose or can other sites on the skin surface serve this purpose? Which body site is the most common source for transmission between carriers or between carriers and fomites? Where on the body are there more likely to be transitions between colonizing and infecting *S. aureus*? Given that there are mutations that adapt the bacterium to sterile site survival, are there colonization sites on the body where these genetic changes have lower fitness cost? Future research can address these questions, supporting the development of interventions to prevent *S. aureus* infections in susceptible people.

## Conclusions

Using genome sequencing we were able to plausibly define USA300 *S. aureus* ISLs as having < 30 SNPs compared to the III. We showed colonization of USA300 after an index USA300 MRSA infection of a patient for up to a year. Persistently colonizing strains could be found in all anatomic sites tested: nose, throat and perirectal area. These results confirm long term persistent colonization by USA300, including MSSA and MRSA, declining over time, and suggest that sites other than the nose can maintain and spread the bacterium around the human body. We estimated that the ISL was introduced to the host’s body not long, approximately 18 weeks, before the index infection. This preliminary study suggests that there would be great value in repeating this design looking at *S. aureus* over more body sites over longer time periods, but at shorter intervals, as well as examining the cloud of diversity of an ISL over time. With this information, it may be possible to define evolutionary constraints giving rise to bottlenecks emerging after exposure to antimicrobials and to determine the natural history of asymptomatic colonization of people by pathogenic *S. aureus* strain types as well as perhaps their risk of recurrent infection.

**List of Abbreviations**
ACME: arginine catabolic mobile element
III: index infection isolate
ISL: intra-subject lineage
MLST: multilocus sequence type
MRCA: most recent common ancestor
MRSA: methicillin resistant *S. aureus*
MSSA: methicillin sensitive *S. aureus*
PVL: Panton-Valentine leukocidin
SSTI: skin and soft tissue infection

## Declarations

### Ethics Approval and Consent to Participate

The study was approved by the Institutional Review Board of the Division of the Biological Sciences of the University of Chicago (protocol 11-0222) and written informed consent was obtained from each subject.

### Availability of Data and Materials

The datasets generated and/or analysed during the current study are available in the NCBI SRA repository under bioproject accession http://www.ncbi.nlm.nih.gov/bioproject/PRJNA388087. An interactive tree with metadata is available at http://itol.embl.de/tree/1701401041483411505410590.

### Authors contributions

TDR performed bioinformatics analysis and wrote the manuscript. RAP performed bioinformatic analysis. ZY processed cultures and performed MLST and PCR genotyping studies on *S. aureus* isolates. TM collected, tested for hemolysis, and froze infecting isolates. MCM recorded much of the clinical data from the medical records. MZD designed and directed the clinical portions of the study and edited the manuscript.

## Acknowledgments

We would like to thank Michael Frisch for help formatting the sequence data for NIH submission.

## Additional Files

**File name:** Additional_File_1.xlsx

**File format:** Excel (.xlsx)

**Title:** Patient metadata

**Description:** Breakdown of all metadata fields collected and summary of genetic results.

**File name:** Additional_File_2.pdf

**File format:** PDF

**Title:** Additional figures and tables

**Description:** Supplementary Table 1: agr frameshifts. Supplementary Figure 1: SNP frequency by genome. Supplementary Figure 2: within ISL estimates for pangenome. Supplementary Figure 3: prophage distribution. Supplementary Figure 4: plasmid distribution. Supplementary Figure 5: variant regions.

